# GAIT BIOMECHANICS OF RABBITS WITH EITHER ACHILLES OR TIBIALIS CRANIALIS ARTIFICIAL TENDONS

**DOI:** 10.1101/2022.01.25.477740

**Authors:** Patrick T. Hall, Caleb Stubbs, Alisha P. Pedersen, Caroline Billings, Stacy M. Stephenson, Cheryl B. Greenacre, David E. Anderson, Dustin L. Crouch

**Affiliations:** University of Tennessee, Knoxville; University of Tennessee College of Veterinary Medicine; University of Tennessee Medical Center, Knoxville; The University of Tennessee Knoxville College of Veterinary Medicine

## Abstract

Artificial tendons have been developed as a replacement for biological tendons with irreparable pathologies and defects. Previous studies reported the mechanical strength and tissue integration of a polyester suture-based artificial tendon, but not its effect on locomotor function. The objective of this study was to quantify the hindlimb biomechanics during hopping gait of New Zealand White rabbits with surgical replacement of either the Achilles (n=2) or tibialis cranialis (TC, n=2) biological tendons with artificial tendons. Once pre-surgery and for five consecutive weeks post-surgery (starting at about two weeks post-surgery), we measured hindlimb kinematics and ground contact pressures with a video camera and pressure mat, respectively. Promisingly, post-surgical locomotor function was either consistent or improved over time in both tendon replacement groups. However, Achilles rabbits exhibited greater immediate post-surgery functional decline and less post-surgical functional recovery than TC rabbits. Compared to healthy rabbits, at the study endpoint, (1) TC rabbits had a 17.3-degree higher (i.e., more plantarflexed) ankle angle at foot strike; and (2) Achilles rabbits had a 39.2-degree lower (i.e., more dorsiflexed) ankle angle at toe off. These functional deficits suggest that the muscles attached to the artificial tendons had lower force-generating capacity. Future studies of artificial tendons are needed to quantify long-term function, determine the effectiveness of structured rehabilitation exercises, and refine surgical implementation.

## INTRODUCTION

Biological tendons, the anatomical links between muscles and bones, transmit forces and store and release elastic potential energy. There are several clinical conditions associated with tendon pathology or defects. Acute tendon rupture, one of the most common tendon injuries^1^, occurs frequently in the Achilles^1-3^, patellar^4^, and quadriceps^5^ tendons. Tendinosis, which involves the breakdown of collagen from chronic overuse^6^, is common in the elbow^7,8^, shoulder^9,10^, and ankle^11^ joints. Rotator cuff tears are most prevalent in older adults but may occur in younger patients following trauma^12^. Limb amputation may damage or remove part or all of the tendons that were attached to the missing limb^13,14^. If the pathology or defect is too severe, the biological tendon may not recover even with clinical treatment, resulting in chronic motor impairment.

One potential treatment is to replace part or all of a damaged or missing tendon with an artificial tendon, defined here as a tendon fabricated from synthetic materials. Developed and tested since as early as 1900, artificial tendons have been made from silk, tantalum wire, nylon monofilament, woven Teflon, polyester, and carbon fiber^15^. More recently, Melvin, *et. al* developed an artificial tendon consisting of braided strands of polyester microfiber suture^16-18^. The advantage of the suture-based artificial tendon is that it can be easily and robustly attached to bone via an orthopedic implant or anchor. Previous studies tested the suture-based tendon *in vivo* (rabbits^17^ and goats^16,18^) and demonstrated that the resulting muscle-tendon junction is stronger than the muscle itself ^17^. Therefore, artificial tendons may be a viable clinical option when tendon repair or other replacement methods (e.g., grafting) are not indicated.

The effect of artificial tendons on motor function is poorly understood but needs to be characterized to support clinical use. Previous *in vivo* studies of the suture-based artificial tendon noted only that animals regained “complete weight-bearing in the operated leg and normal gait by 3 [weeks post-surgery]” with no supporting data^18^. Based on well-known musculoskeletal mathematical relationships^19^, we expect that the effect of an artificial tendon on motor function will depend on several factors. For example, the mechanical properties (e.g., stiffness) of the artificial tendon should influence not only how it transmits force and stores energy, but also the attached muscle’s excursion, mechanics, and energetics^20^. How the artificial tendon is attached to muscle may change how forces are exchanged between them or affect the health and structure of the muscle. The length of the artificial tendon will influence the muscle’s force generating capacity over a given joint range of motion^21^. Finally, formation of adhesions on the surface of the artificial tendon, as on other implants^22^, may restrict or prevent tendon sliding and force transmission.

The goal of this study was to quantify the effect of a suture-based artificial tendon on locomotor function in a rabbit model. Once pre-surgery and five consecutive weeks post-surgery, we quantified sagittal-plane kinematics and ground contact pressures in New Zealand White rabbits that had either the biological Achilles tendon (n=2; A1 & A2) or tibialis cranialis insertion tendon (n=2, TC1 & TC2) surgically replaced with an artificial tendon. We hypothesized that locomotor function would decrease immediately following surgery but subsequently recover toward pre-surgical levels.

## METHODS

### Animal Model

All animal procedures were approved by the University of Tennessee, Knoxville Institutional Animal Care and Use Committee. We tested artificial tendons in New Zealand White rabbits since they are a small mammal but large enough to test physical prototypes of suture-based artificial tendons. Additionally, we observed in a previous study^23^ that rabbits have a nearly plantigrade hopping gait and, thus, have a large ankle range of motion compared to other similar-sized mammals that have a digitigrade gait (e.g., dogs, cats); the large range of motion makes it easier to distinguish differences in kinematics among experimental groups.

### Artificial Tendon

The artificial tendons used in this study were based on the prostheses developed by Melvin, *et al*. ^16,18^, except that we used *braided* rather than *unbraided* polyester microfiber suture. Specifically, we used customized USP size 0 braided polyester suture cut to 12” length and double-armed with swaged 3/8-circle taper point needles (0.028” wire diameter) (RK Manufacturing Corp, Danbury, CT, USA). The sutures were grouped into bundles of 2 strands for the tibialis cranialis tendon and 3 strands for the Achilles tendon. The suture bundles were folded in half and braided, placing a loop at the distal end and the needles at the proximal end (Fig. 1). The artificial tendons were coated in biocompatible silicone (BIO LSR M340, Elkem Silicones, Lyon, FR) over the braided section to discourage buildup of fibrotic adhesions. The tendons were cleaned and sterilized before surgery.

**Figure 1.**
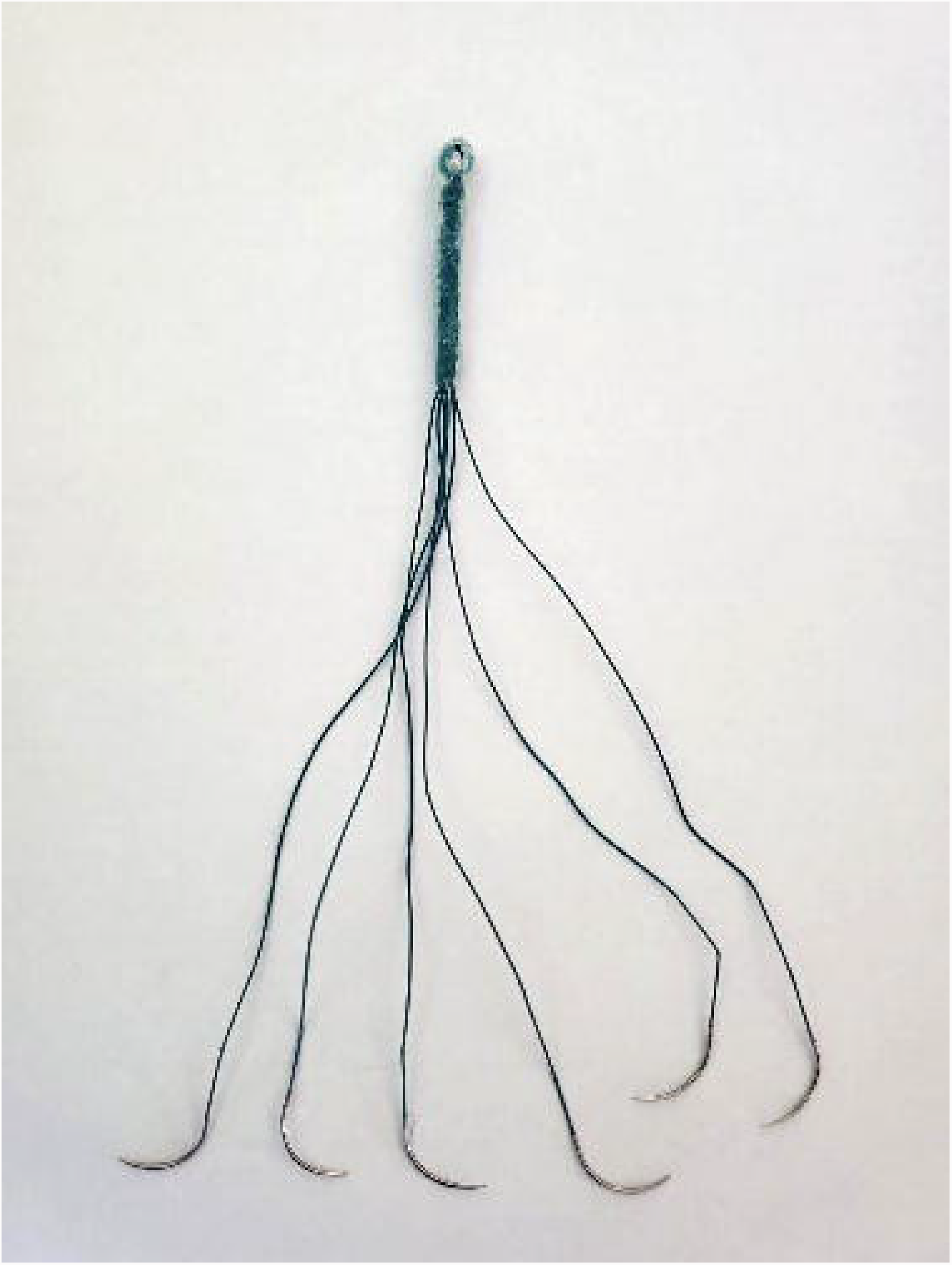
Example of the suture-based artificial tendon used to replace the Achilles tendon.

### Surgical Procedure

Four rabbits (16 weeks old, 3.40 ± 0.26 kg at the time of surgery) were divided into two groups to undergo surgical replacement of either the Achilles (A1 & A2) or Tibialis Cranialis (TC1 & TC2) biological tendon with an artificial tendon. Under anesthesia, the respective biological tendon was excised from the muscle-tendon junction to the bony insertion point. The artificial tendon was sewn into the proximal end of the muscle as previously described^16^. We attached the artificial tendon to the bone using a suture anchor into either the superior aspect of the calcaneus (Achilles) or the lateral aspect of the talus (tibialis cranialis).

### Post-Surgical Care

The limb was bandaged starting immediately post-surgery until about two weeks post-surgery to protect the incision site and prevent overloading the muscles and artificial tendons during the early integration period. To monitor the integrity of the artificial tendon *in vivo*, we performed radiography of the operated limb every other week, starting on the day of surgery. When analyzing the radiographs, we inspected for potential mechanical failure of either 1) the suture connecting the artificial tendon and suture anchor, 2) the suture anchor fixation in the bone, or 3) the muscle-tendon interface. After bandages were removed, the rabbits were given pen time to promote limb use.

### Biomechanics Testing

We quantified biomechanics during hopping gait non-invasively. A pressure mat (Tekscan, Very HR Walkway 4; South Boston, MA) was placed inside an acrylic tunnel, and a webcam (1080P HD Webcam, SVPRO) was placed approximately 3 feet to one side of the acrylic tunnel to record sagittal-plane kinematics. The pressure mat and video data were synchronously recorded at 60 Hz through Tekscan Walkway software. Before testing, we shaved both hindlimbs of each rabbit and marked the approximate joint centers of the knee, ankle, and metatarsophalangeal (MTP) joints with black ink, based on bony landmarks.

For each trial, the rabbits were prodded to hop along the pressure mat through the acrylic tunnel. The width of our pressure mat (11.2cm) permitted only unilateral pressure recording. Therefore, each rabbit completed about 10 trials of hopping in each direction through the tunnel while we recorded biomechanics data for the hindlimb closest to the camera. Each trial began when the rabbit entered the tunnel, and a trial was deemed successful if the rabbit continued the hopping motion through the entire length of the tunnel without stopping. Each rabbit underwent 6 biomechanics testing sessions (S0-S5) over the course of the study. We conducted the first testing session during the week before surgery (S0). After surgery, we waited about 2 weeks for the rabbits to recover before starting weekly post-surgical testing for five consecutive weeks (S1-S5).

### Data Processing and Statistical Analysis

We processed and analyzed data only for the stance phase of hopping gait, when the hindlimb experiences the most biomechanical loading due to contact with the ground. We calculated the hindlimb kinematics from the frames of the synchronized video corresponding to stance phase. Frame-by-frame, a custom MATLAB (Mathworks, Natick, MA) program (1) identified the centroids of joint centers marked with black ink, (2) defined foot and shank limb segments as lines connecting adjacent joint centers, and (3) calculated the foot and ankle joint angles from the limb segment lines (Fig. 2).

**Figure 2.**
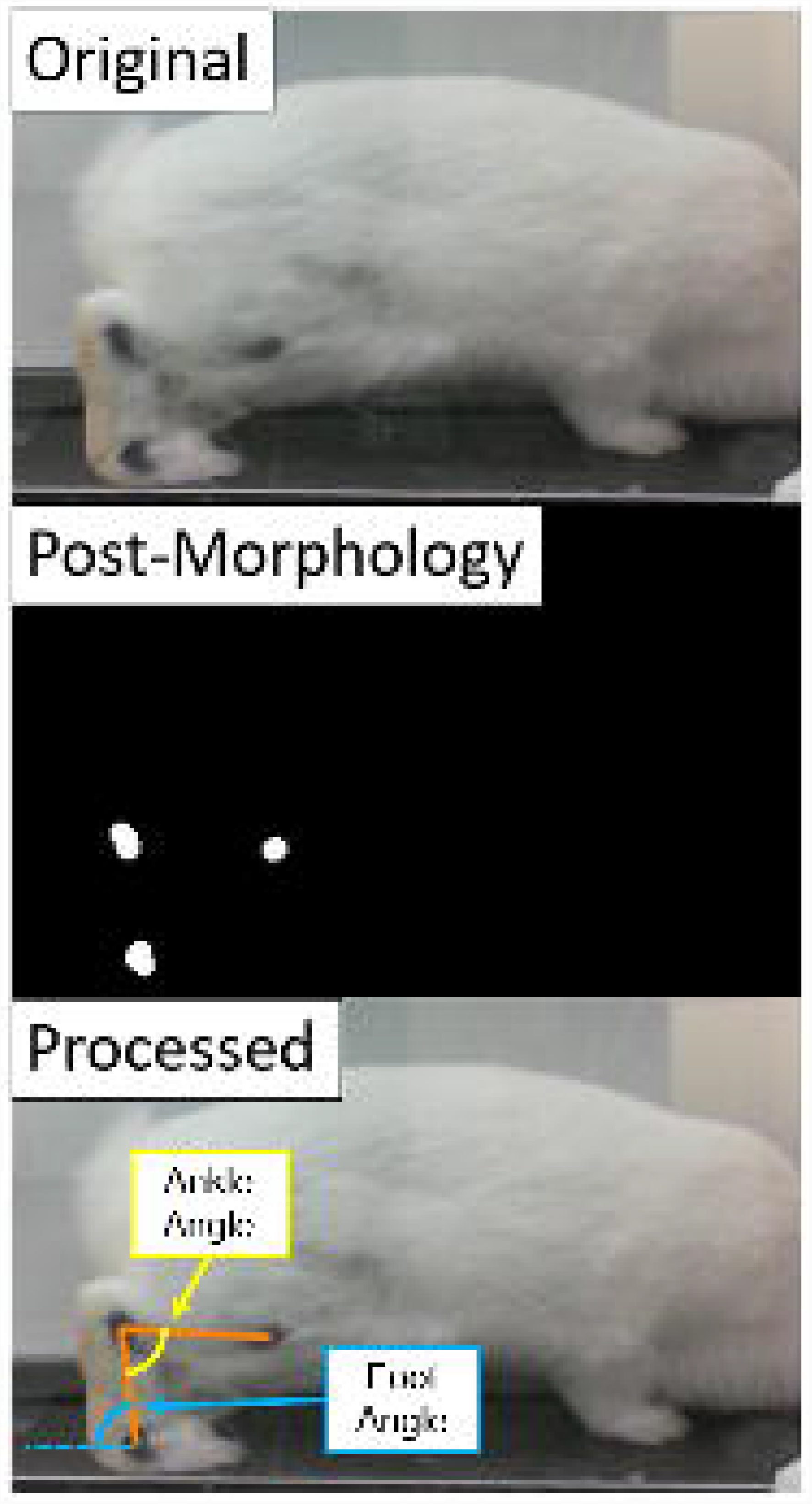
Method used to measure hindlimb kinematics in the sagittal plane from videos. Limb segments (orange) were defined between the centroids of the joint markers. We calculated the ankle angle (yellow) as the angle between the shank and foot segments; and the foot angle (blue) as the angle between the foot segment and ground.

Analyzing joint angles individually masks changes that occur in the whole kinematic chain of the hindlimb. Therefore, we computed the position of the knee joint center in the sagittal plane throughout stance using the foot and ankle angles and a generic planar kinematic model of the rabbit hindlimb ^23^. The foot and shank segments of the kinematic model were 2 and 3 normalized length units (NLUs), respectively, based on the approximate anatomical length proportions averaged across all rabbits.

To analyze the pressure data, in the Tekscan Walkway software, we first isolated the contact foot of the hindlimb of interest by drawing a strike box around the area of contact. At each timepoint, we computed the average ground contact pressure, contact area, and vertical ground reaction force (vGRF) from the pressure values within the strike box. vGRF was expressed as a percentage of body weight (%BW). We computed the duration of stance phase as the length of time over which the total ground contact pressure within the strike box was greater than zero.

We evaluated the extent of post-surgery recovery by comparing *time-series biomechanical data* (foot and ankle angle, vGRF, contact area, pressure, and knee joint center position) between each experimental group (n=2 per group) at S5 and a control group of healthy rabbits (n=6) measured in a previous study^23^. We compared normalized knee joint center positions between groups by calculating the Euclidean distance between positions at each timepoint. To permit comparison among trials of different stance durations, for each trial, we normalized the timepoints of the stance phase data by the duration of stance phase. The normalized time-series data were averaged across trials and rabbits in each group and compared between experimental and control groups using a non-parametric Mann-Whitney U-Test at every percent of stance with α=0.05 ^24^.

We compared *summary outcome measures* to identify functional differences (1) between pre- and post-surgery testing sessions and (2) across post-surgery testing sessions. The summary outcome measures, computed from the time-series data for each trial, were: (1) range of motion (difference between the maximum and minimum joint angle during stance) for each joint, (2) the minimum and maximum joint angles during stance, (3) the joint angles at foot strike and toe off, and (4) the average vGRF, contact area, and pressure across stance. For each testing session, the summary outcome measures were averaged across trials for each rabbit, then averaged across rabbits within each experimental group. Due to our small sample size, we compared summary outcome measures qualitatively to identify trends, rather than qualitatively using a statistical test (e.g., ANOVA).

## RESULTS

### Time-Series Outcomes at the Final Timepoint (S5)

#### Tibialis Cranialis (TC) Group

At S5, foot and ankle angles were significantly different between the TC and control groups during the first 35% of stance (Fig. 3); however, the magnitude of the difference during this period was small (mean difference = 10.45 degrees). The minimum ankle angle occurred at about 40% of stance phase for both TC and control groups.

**Figure 3.**
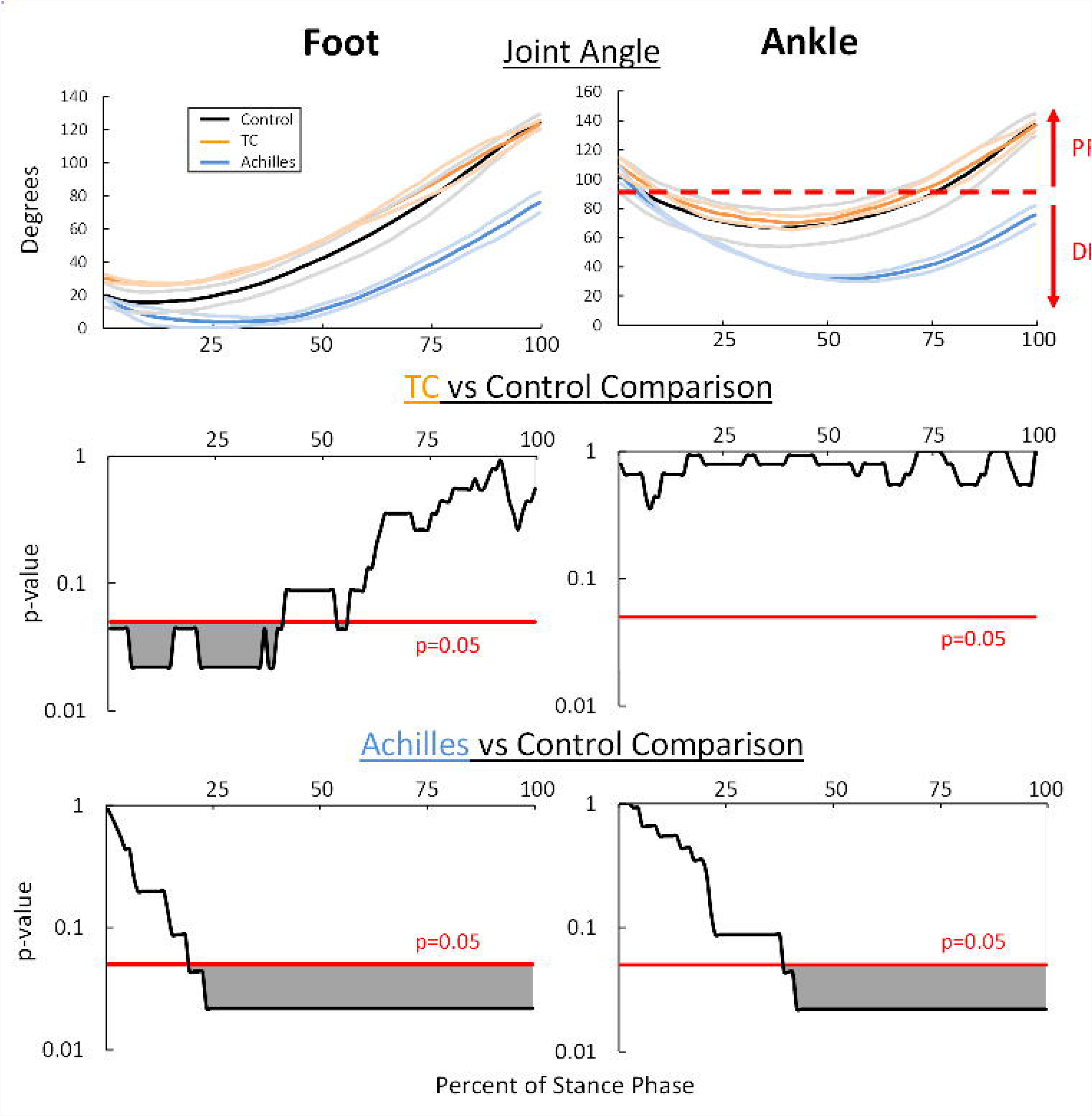
(top row) Time-series joint angles during stance phase (top row) and (middle and bottom rows) results of Mann-Whitney U-Test to compare between experimental and control groups. Compared to the control group, joint angles were similar in the TC group but significantly lower in the Achilles group during most of stance. Groups were significantly different where p>0.05 (grey shaded regions). PF=plantarflexion, DF=dorsiflexion.

During stance phase, rabbits with TC replacement at S5 and the control group had similar normalized knee joint center kinematics (Fig. 4). At foot strike, the foot angle was higher in the TC group than in the control group, resulting in a maximum distance of about 0.4 NLUs between each group’s knee joint centers. As the rabbits progressed through stance phase, the knee joint center kinematics of the TC and control groups became more similar, with distance of 0.05 NLUs between groups at toe off. The distance between TC and control groups averaged across stance was 0.17 NLUs.

**Figure 4.**
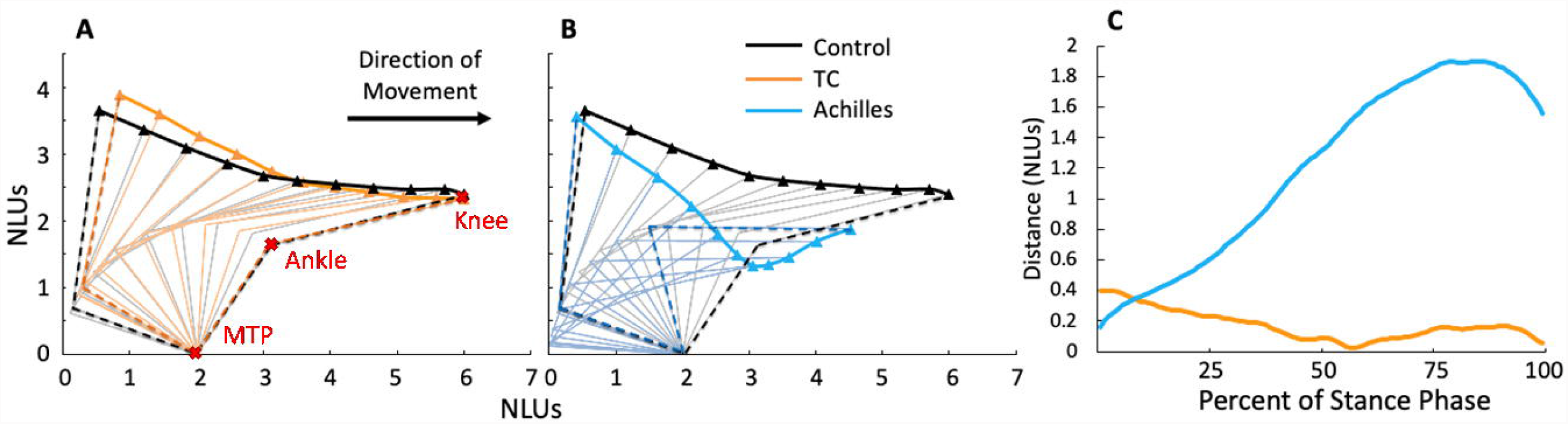
**(A, B) Normalized knee joint center positions during stance phase at S5**. Compared to the control group, the progression of knee joint center position over the stance phase was similar in the TC group but lower and less forward in the Achilles group. **(C) Distance in knee joint center position between each experimental group and the control group. For the TC group, the d**istance was relatively small (within 0.4 NLUs) and constant. For the Achilles group, the distance started small but increased to a much higher value (nearly 2 NLUs) by the end of stance.

The average ground contact pressure was significantly different between the TC group at S5 and the control group during 18-89% of stance phase (Fig 5). Both contact area and vGRF were not significantly different between TC and control groups throughout most of stance.

**Figure 5.**
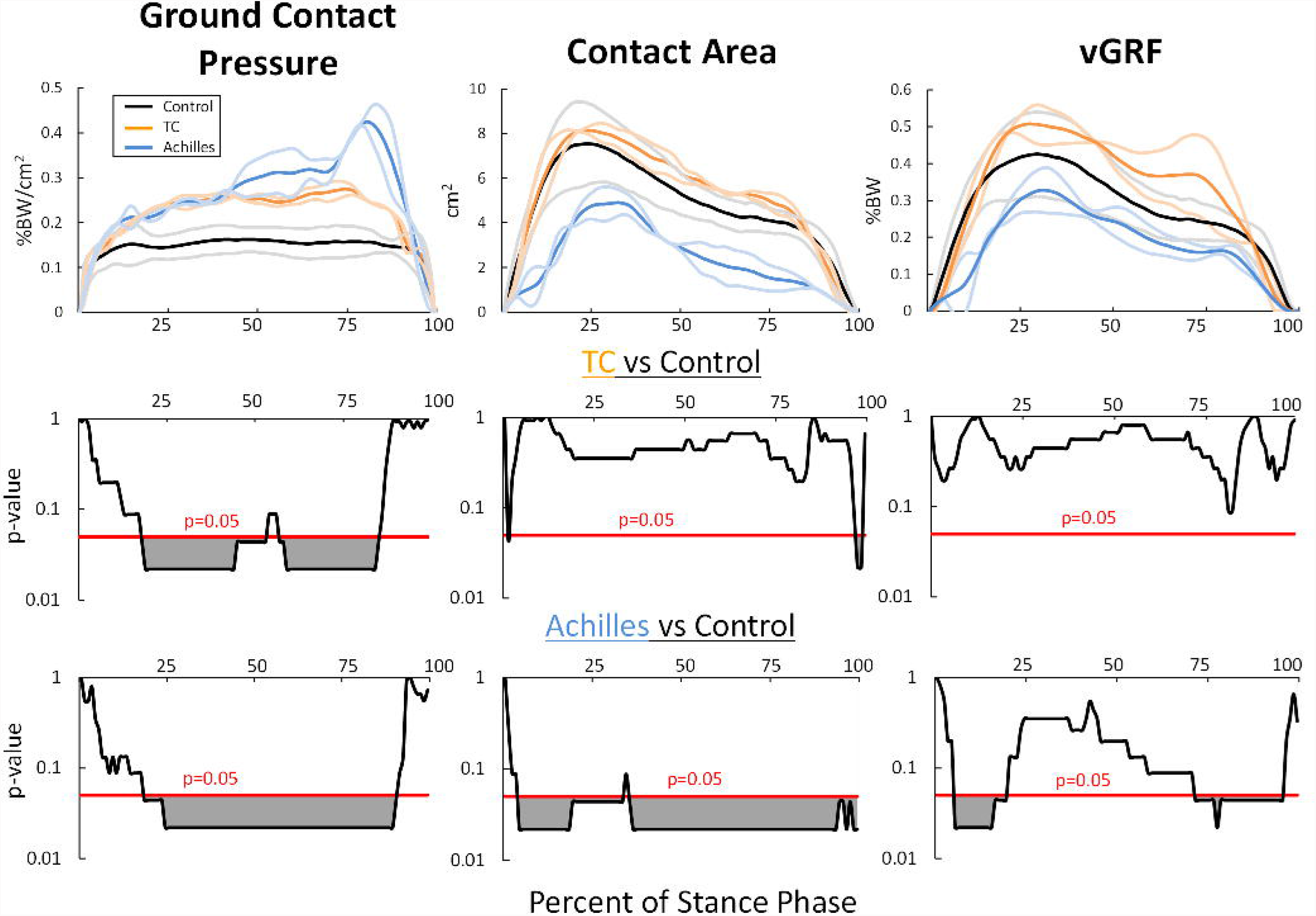
(top row) Average ground contact pressure, contact area, and vertical ground reaction force (vGRF) during stance phase at S5. (middle and bottom rows) Results of Mann-Whitney U-Test to compare between experimental and control groups. Lighter lines represent ± standard deviation. Ground contact pressure was significantly higher in the TC and Achilles groups than in the control group during 20-90% of stance. Compared to the control group, contact area and vGRF were similar in the TC group but significantly lower in the Achilles group for much of stance. Groups were significantly different where p>0.05 (grey shaded regions).

#### Achilles Group

At S5, rabbits with Achilles replacement had significantly different foot and ankle angles starting at 15% and 31% of stance phase, respectively, compared to the control group (Fig 3). The minimum ankle angle was 34.4° lower and occurred later in stance (57%) in the Achilles group than in the control group (40%). Compared to the control group, both foot and ankle angles in the Achilles group were similar at foot strike but became and stayed lower throughout most of stance. Consequently, the knee joint center position dipped lower and had less forward progression in the Achilles group than in the control group (Fig. 4). The maximum distance between the knee positions of the Achilles and control groups was 1.89 NLUs, which occurred at 87% of stance. The average distance between groups across stance was 1.2 NLUs.

Compared to the control group, the Achilles group had significantly different (qualitatively lower) vGRF during 7-20% and 72-89% of stance (Fig. 5) and significantly different (qualitatively lower) contact area during 6-22% and 39-94% of stance. However, the lower contact area resulted in up to three times higher pressure during 19-92% of stance phase compared to the control group.

### Longitudinal Summary Outcomes

#### TC Group

Foot and ankle range of motion (ROM) were similar across all test sessions (Fig. 6). The foot angle at foot strike increased from 11.5 degrees at S0 to 28.8 degrees at S1 and maintained this lack of dorsiflexion at foot strike in subsequent sessions (Fig. 7). However, the ankle angle at foot strike and the foot and ankle angles at toe off were similar across all pre- and post-surgery test sessions. The vGRF and contact area, averaged across stance, decreased from S0 to S1 by 6 %BW and 1.22 cm^2^, respectively, but returned to pre-surgical levels by S3 (Fig. 8). The average ground contact pressure values were similar across test sessions.

**Figure 6.**
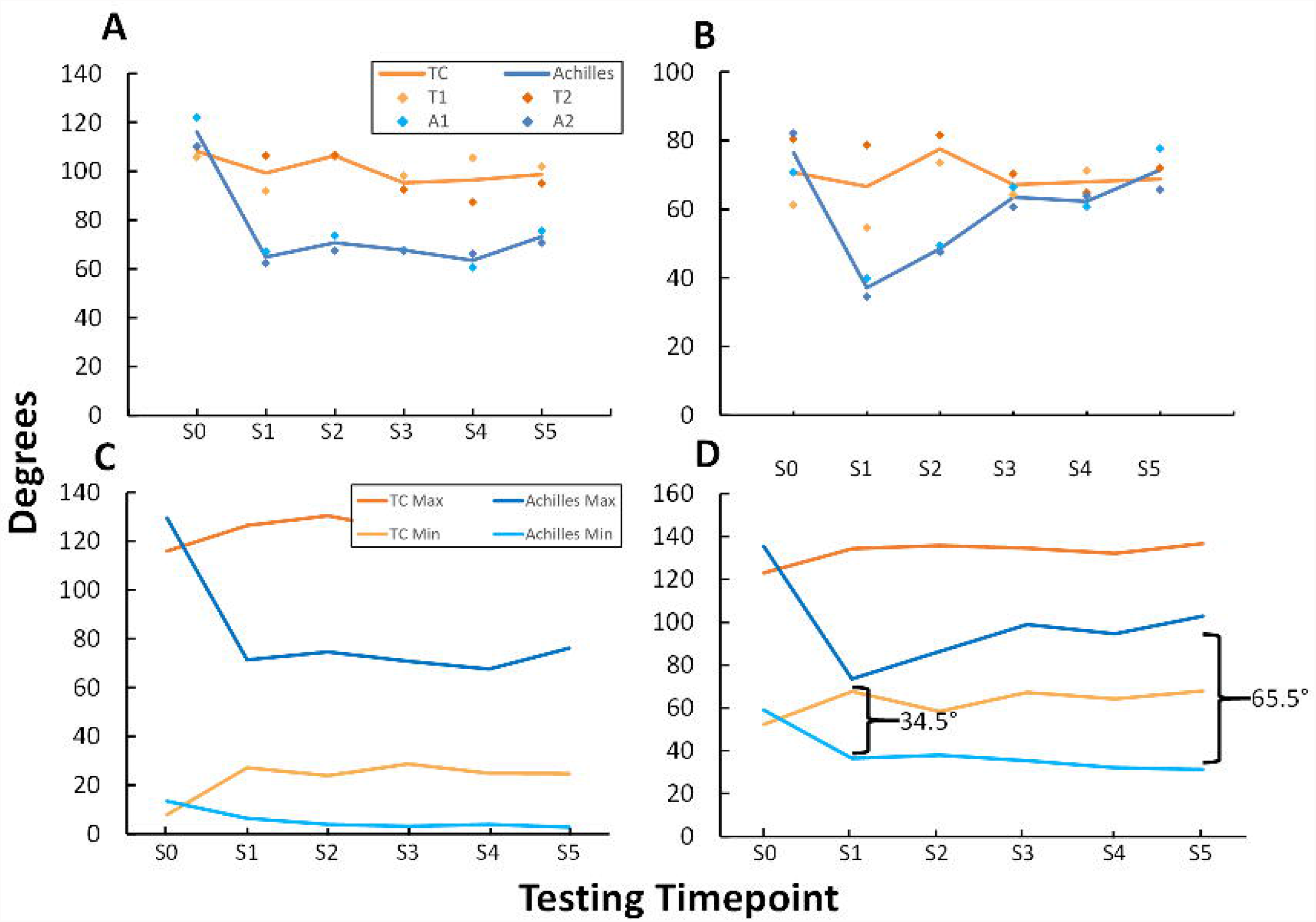
**(A) Foot angle and (B) Ankle angle range of motion (ROM) at pre- (S0) and post-surgery (S1-5) testing sessions**. The lines represent the average of the two samples in each group at each session. The TC group had similar ROM across all sessions. In the Achilles group, ROM decreased substantially from S0 to S1; from S1 to S5 ankle angle ROM but not foot angle ROM increased to pre-surgical levels. **(C) Minimum and maximum Foot and (D) Ankle angles during stance**. The increase in ankle angle ROM in the Achilles group was due to an increase in the maximum angle rather than a decrease in the minimum angle.

**Figure 7.**
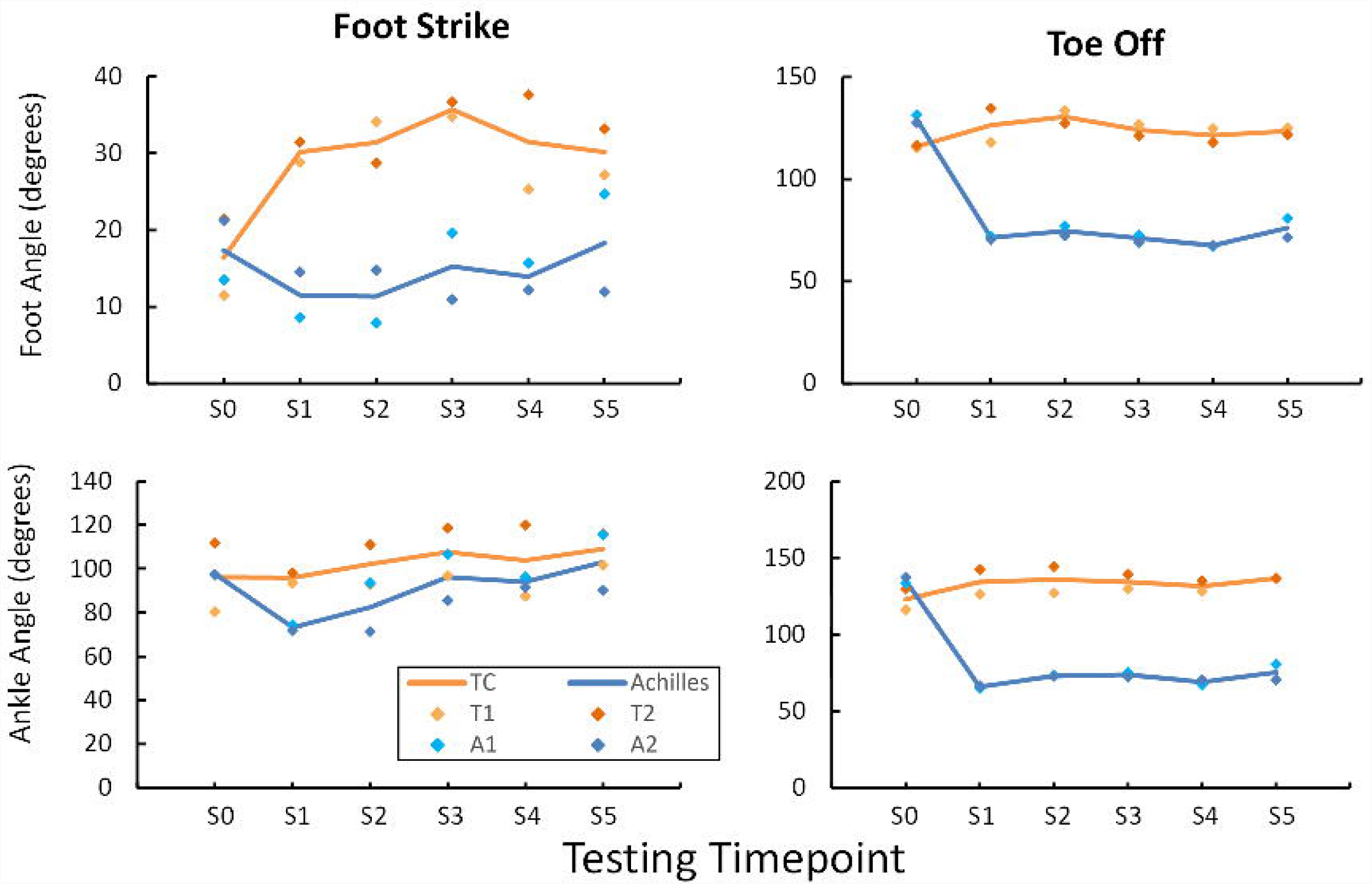
Joint angles at foot strike (left) and toe off (left) at pre-surgery (S0) and post-surgery (S1-5) testing sessions. The lines represent the average of the two samples in each group at each session. At foot strike, the TC group had similar ankle angles across all sessions but higher foot angle post-surgery that never recovered. In the Achilles group, foot and ankle angles at foot strike decreased slightly from S0 to S1 but recovered by about S3. At toe off, joint angles in the TC group were similar across all sessions but decreased substantially from S0 to S1 and never recovered in the Achilles group.

**Figure 8.**
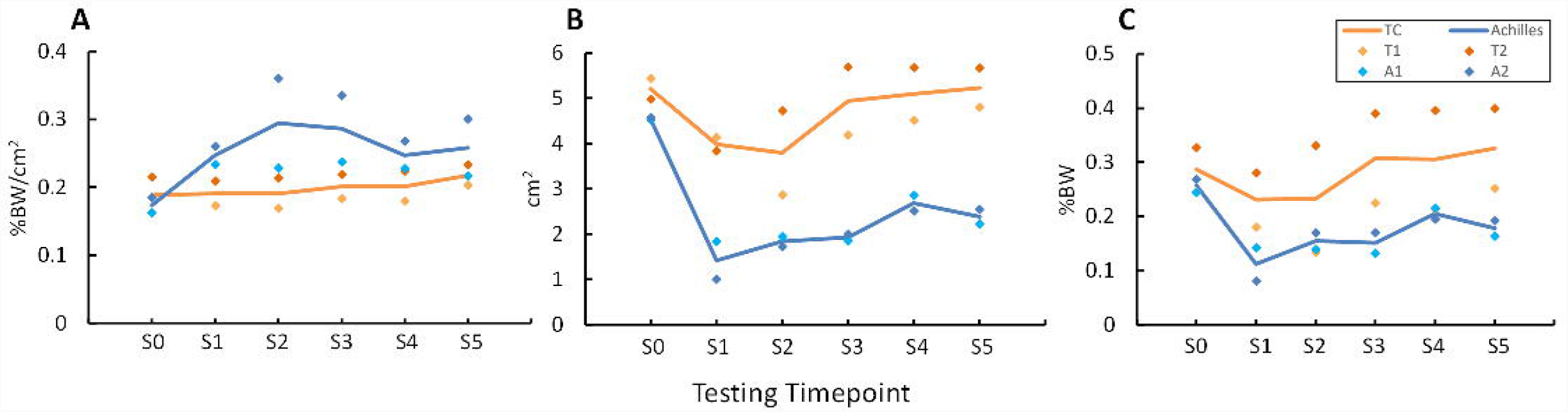
**(A) Ground contact pressure, (B) contact area, and (C) vertical ground reaction force (vGRF) at pre-surgery (S0) and post-surgery (S1-5) testing sessions**. In the TC group, pressure was the same across all sessions, while contact area and vGRF decreased slightly from S0 to S1 but recovered by about S3. In the Achilles group, pressure increased from S0 to S1 but recovered toward pre-surgery levels starting at S3; contact area and vGRF decreased from S0 to S1 but steadily increased toward pre-surgery levels from S1 to S5.

#### Achilles Group

Foot and ankle angle ROM decreased by 51.3 and 39.2 degrees, respectively, from S0 to S1 (Fig. 6). The foot angle ROM did not increase post-surgery. However, ankle angle ROM increased to pre-surgical levels by S5. The increase in ankle angle ROM between S1 and S5 was primarily due to an increase in the maximum ankle angle toward greater plantarflexion (S1: 73.5 degrees; S5: 102.9 degrees), rather than a decrease in the minimum ankle angle toward greater dorsiflexion (S1: 36.4 degrees; S5: 31.3 degrees). At foot strike, both the foot and ankle angles were similar across test sessions (Fig. 7). However, the foot and ankle angles at toe off decreased by 58.3 and 69.6 degrees, respectively, from S0 to S1 and never increased toward pre-surgical levels. The average vGRF and contact area during stance decreased by 14.5 %BW and 3.13 cm^2^, respectfully, from S0 to S1 but increased by 6.6 %BW and 0.96 cm^2^, respectively, toward pre-surgical values from S1 to S5 (Fig. 8). The average ground contact pressure during stance increased from S0 to S1, but at S3 began decreasing back toward pre-surgical levels.

## DISCUSSION

In support of our hypothesis, our data suggest that rabbits with either tendon replacement (Achilles or TC) recovered toward pre-surgical (S0) levels of function for at least part of the biomechanical variables we measured. By S5, the TC group was closer to S0 function levels than the Achilles group. However, there was also less functional decline from S0 to S1 and, thus, less function to recover to achieve S0 function levels in the TC group than in the Achilles group. One potential explanation for the difference between groups is a difference in muscle-tendon loading between tendons during stance. Ankle plantarflexion torque is expected to be greater than dorsiflexion torque during stance since plantarflexion contributes to body weight support and forward propulsion that occur during stance. The triceps surae muscles (gastrocnemius and soleus), which are attached across the ankle by the Achilles tendon, are considered ankle plantarflexors in rabbits^25^. There may have also been a difference in how the triceps surae and TC muscles were affected by the tendon replacement as discussed below.

The functional deficits in the Achilles group, compared to healthy rabbits, included lower (i.e., more dorsiflexed) ankle angle and vGRF. These two deficits reflect a weakness in ankle plantarflexion. Simulations of ankle plantarflexor weakness in humans similarly predicted a more dorsiflexed ankle throughout most or all of stance^26^. Thus, the more dorsiflexed posture in the Achilles group suggests that the triceps surae muscle group generated less force with the artificial tendon than with the biological tendon.

The only statistically significant difference in time-series hindlimb kinematics between the TC group at S5 and healthy rabbits was a higher (i.e., more plantarflexed) foot angle during the first 35% of stance phase in the TC group. A higher foot angle during early stance resembles a “drop foot” condition. In humans, drop foot is a result of insufficient (i.e., low) ankle dorsiflexion joint moment and, consequently, dorsiflexion angle during swing. Drop foot presents with several neurological and musculoskeletal conditions, including stroke^27^ and strain or tear of the Tibialis Anterior tendon ^28,29^. The TC muscle is considered an ankle dorsiflexor in rabbits^25^. Therefore, the observed higher foot angle during early stance and its resemblance to drop foot suggest that the TC muscle generated less force with the artificial tendon than with the biological tendon.

The muscles may have produced less force with artificial tendons for several reasons. For one, the length of the artificial tendon could have been longer or shorter than the replaced biological tendon, which would shift the muscle toward the ascending limb (shorter muscle) or descending limb (longer muscle), respectively, of the muscle force-length curve^19^. Depending on the normal operating range of the muscle on the force-length curve, the shift may decrease the maximum force the muscle can produce over a given joint range of motion. The muscles attached to the artificial tendons may have also had lower inherent force generating capacity for a few possible reasons. For example, implanting the suture-based artificial tendon may have caused minor muscle damage, though previous histology showed good integration between the suture and muscle with only mild capsular fibrosis^30^. The muscle may have also atrophied with decreased use^31^ due to bandaging (i.e., immobilization) and discomfort following surgery.

It is well known that the body can form a fibrous, collagenous capsule around implanted devices to isolate it from the surrounding tissues. The fibrotic capsule could cause functional complications for devices, like the artificial tendon, that need to move relative to surrounding tissues to function properly. In our study, upon *ex vivo* dissection of one specimen (A2), we observed fibrotic tissue fusing the artificial tendon to the flexor digitorum insertion tendon before it crossed the ankle. Pulling either the triceps surae (Achilles) or flexor digitorum muscle groups individually caused co-flexion of the ankle and digits. This is functionally undesirable and precludes independent movement at the two joints. For example, the fused tendons likely explain our qualitative observations, from motion videos, of excessive plantarflexion of the digits in the operated limb, which prevented the mid- and hindfoot from contacting the ground and resulted in lower contact area during stance (Fig. 7). Thus, to maximize motor function, new approaches are needed to prevent or reduce the fibrotic capsule on artificial tendons.

This study had several limitations. First, though we had multiple tendon sizes available (in 2 mm increments) during surgery, we did not have a way to adjust the length of the artificial tendon after implantation. Being able to adjust tendon length intraoperatively would address the functional complications of having an improper tendon length, described above. Second, our sample size was small, with only 2 rabbits per experimental group. This limited the power of our time-series statistical comparisons, and we chose not to perform a statistical analysis of summary outcome measures. However, both rabbits in each group showed similar trends that supported our hypothesis. Third, during post-surgical recovery, the rabbits were not given formal, structured physical therapy, which may have limited the rate and extent of functional recovery the rabbits could have achieved during the study. Daily enrichment and pen time may have had some therapeutic effect. Fourth, we had a relatively short post-surgical testing period. Though the TC group fully recovered stance phase biomechanics according to our outcome measures, the Achilles group may have recovered more with more time. Fifth, due to technical limitations of our testing setup, we measured biomechanics from one limb at a time and during stance phase only. In future studies, we plan to use a larger pressure mat and capture motion at a higher frame rate to permit simultaneous bilateral data collection of both stance and swing phases of gait.

In conclusion, our preliminary results showed, promisingly, that rabbits with an artificial tendon either preserved or recovered locomotor function. The Achilles group experienced greater post-surgical functional decline and had poorer function at the study endpoint than the TC group. In future studies, we plan to measure function for longer duration and implement formal rehabilitation to quantify the maximum possible extent of recovery. Our results can inform the development and clinical translation of artificial tendons for patients with irreparable tendon pathologies and defects.

## ACKNOWLEDGMENTS

The authors thank Dr. Bryce Burton, Dr. Kelsey Finnie, Dr. Lori Cole, and Chris Carter for veterinary care provided for the rabbits in this study, and the Office of Laboratory Animal Care and Animal Housing Facility staffs at the University of Tennessee, Knoxville for animal care assistance. Thank you to Elizabeth Croy for her assistance in surgical prep and operation. Thanks to Dr. Katrina Easton for providing constructive feedback on the manuscript. Research reported in this publication was supported by (1) the Eunice Kennedy Shiver National Institute of Child Health & Human Development of the National Institutes of Health under Award Number K12HD073945, (2) NSF CAREER Award #1944001, (3) a seed grant from the University of Tennessee Office of Research and Engagement, and (4) the University of Tennessee Department of Mechanical, Aerospace and Biomedical Engineering start-up funds.

